# Activity throughout the lichen phylogeny indicates a focus on regulation of specialized metabolites

**DOI:** 10.1101/2020.07.09.195743

**Authors:** Ludmila V. Roze, Maris Laivenieks, Kristi Gdanetz, John E. Linz, Alan M. Fryday, Frances Trail

**Affiliations:** Department of Plant Biology, Michigan State University, East Lansing, MI 48824, USA; Department of Internal Medicine, University of Michigan, Ann Arbor MI 48109, USA; Department of Food Science and Human Nutrition, Michigan State University, East Lansing, MI 48824, USA; Department of Plant, Soil and Microbial Sciences. Michigan State University, East Lansing, MI 48824, USA

**Keywords:** lichens, specialized metabolites, antifungal, microbiome, *Aspergillus*, *Fusarium*, aflatoxin, trichothecenes

## Abstract

Lichens are complex multi-microorganismal communities that have evolved the ability to share their thalli with a variety of microorganisms. As such, the lichenized fungus becomes a scaffold for a variety of microbes and occasionally insects. Lichens are known to produce a plethora of unique specialized (secondary) compounds that demonstrate biological activities, including antibacterial, antifungal, antiviral, and antioxidant, that may provide protection from harmful microbes. The longevity of lichens and their robustness, despite a close association with diverse microbes, provides an interesting study system to view the role of specialized metabolites in managing a microbial community. The objective of this study was to identify the effects lichens may have on basic functions of fungi in and on the lichens. We tested chemical extracts from lichen species across the phylogenetic tree for their effects on sporulation, hyphal growth and specialized metabolite production, using two well-studied mycotoxigenic fungi (*Aspergillus parasiticus* (aflatoxin) and *Fusarium graminearum* (trichothecenes) whose functions are easily observed in culture. By far the most prevalent activity among the 67 lichens we tested were effects on accumulation of fungal specialized metabolites, which appeared in 92% of the lichen species analyzed across the phylogeny, although the lichen extracts were also active against fungal sporulation (31%) and growth (12%). The consistent presence of this regulatory activity for specialized metabolism indicates this is an important aspect of lichen integrity. Interestingly, inhibition of accumulation of products of the aflatoxin biosynthetic pathway was the predominant activity, whereas increased accumulation versus decreased accumulation of the production of trichothecenes were about equal. This suggests multiple mechanisms for addressing fungal processes. We performed microbiome analysis of four lichen species and identified oomycetes as members of the microbiomes. Although a small sample size was used for comparing microbiomes, the lichen species exhibiting lower effects on the test fungi had a higher number of OTUs. Members of the lichen community may manipulate specialized metabolism of the essential and transient fungal members and thus attenuate negative interactions with the incumbent fungi or, alternatively, may support the production of compounds by beneficial fungal partners. The ability to control the microbiome by specialized metabolites as opposed to controlling by reducing sporulation of growth, can be effective, discerning, and energetically thrifty, allowing the microbiome members to be controlled without being invasive. Elucidating the role of specialized metabolites in the mechanisms underlying lichen assembly and function has important implications for understanding not only lichen community assembly but for revealing the fundamental processes in microbiota in general.

## Introduction

The lichen microbiome consists of the principle myco- and photobionts, and numerous other lichenicolous and endolichenic fungi, bacteria, and occasionally insects. The longevity of lichens and their robustness despite a close association with diverse microbes provides an interesting study system to view the role of specialized (secondary) metabolites in managing a microbial community. Lichens are known for their unique specialized metabolites that have been used in many cases to human benefit as antibiotics, antifungals, and antivirals.

The contribution of the photobiont, mycobiont, or other members of the microbial community in generating these metabolites is not well understood. Recently, studies on the lichen microbiome have been published, with particular emphasis on lichen associated fungi. Endolichenic microbes are known to produce unique specialized chemistries that may benefit the lichen as a whole. Recent findings indicated that a major lichen metabolite is strongly correlated with the presence of a basidiomycete yeast and is either produced by it or it induces its production by the mycobiont (Spribille *et al.* 2016). LePogam *et al.* (2016) demonstrated that different specialized metabolites had specific locations within the lichen thallus, implying a specific site of synthesis and providing clues to the organism of synthesis. Grube et al (2015) used metagenomics and metaproteomics to show that bacteria within the lichen microbiome provide nitrogen fixation, some vitamins, and have the capacity to alleviate desiccation and salinity stress (reviewed in Aschenbrenner *et al.* 2016). They also revealed evidence for antagonistic roles to other microbes in the form of antimicrobials. Bacteria may play a role in recycling nutrients from older parts of the thallus (Grube *et al.* 2009, 2012; Hodkinson *et al.* 2012; Aschenbrenner 2015; Calcott *et al.* 2018). Many studies focus on identification of medicinally active compounds, have begun to characterize these metabolites and reveal their roles in lichen function, and have identified novel antimicrobials (Calcott *et al.* Chem 2018; Sierra *et al.* 2020).

Previously, we developed high throughput assays to identify, isolate and characterize compounds that can be used to control mycotoxins in plant pathogens (Annis *et al.* 2000), including compounds from black pepper that specifically affect transcription of genes involved in aflatoxin biosynthesis (Trail & Jones. 2018; 2019). Previous reports have suggested that mycotoxigenic fungi are part of the lichen microbiome and that mycotoxins, including aflatoxins, deoxynivalenol, alternariol, and sterigmatocystin, were detected in lichens and stable for many years in herbarium collections (Burkin & Kononenko 2015; Burkin *et al.* 2012; Ji *et al.* 2016). The function of the mycotoxins, if any, for the lichen community is currently unknown.

We adapted an activity screen of lichen extracts across the lichen phylogeny, with emphasis on the Lecanoromycetes, which are most abundant in our area. The objective of this study was to determine the types of antifungal activities that are prominent in lichen thalli. We tested for three activities that would be important in a host-microbiome relationship: growth, sporulation and production of specialized metabolites. Identification and characterization of these activities would be essential to determining their role, so we developed a pathway to identify and characterize compounds with activities in lichen species across the phylogenetic tree on potential colonizing fungi that test for their effects (Figure S1). Our goal was to initiate a broader understanding of the distribution of activities present, to enhance our knowledge of the complexity of the lichen community through microbiome analysis, and thus to identify the tools lichens and their colonizers have to affect the community.

## Materials and Methods

### Lichens collection and storage

Lichens were either collected fresh or stored herbarium specimens were used. Fresh lichen material was air dried and stored at room temperature in a paper bag. Species authentication was performed based on morphological examination. For isolation of lichenized fungal cultures, fruiting bodies were collected fresh and frozen in 2 ml screw-cap glass tubes at −20 C until used for isolation of ascospores. Photobionts were isolated from fresh lichen thalli. Referenced voucher specimens were stored in the Trail lab and submitted to the Michigan State University Herbarium.

### Preparation of lichen extracts

Individual lichens were dried, cut into 1–4 mm^2^ pieces, and approximately 100–200 mg were extracted twice consecutively in 2 ml solvent (ethanol, acetone, hexane or ethyl acetate) in glass vials. The two extracts were combined and evaporated under N_2_, redissolved in the corresponding solvent (0.25 ml per 100 mg lichen extracted), and stored at –20°C.

### Culturing mycobionts and photobionts

Fungal cultures were grown on malt extract (Gibco™ Bacto™ Malt Extract, Fisher Scientific, Hampton, NH) agar medium (MEA) from isolated ascospores. The photobiont of *C. cristatella* was isolated from ground holobiont by the dilution method on MEA.

### High through-put activity assay of effects of lichen extract on mycotoxin biosynthesis, hyphal growth and sporulation

To facilitate the identification of lichen species that produce bio-active compounds, a high through-put activity assay was developed as a primary visual screen for the effect of lichen extracts on hyphal growth, sporulation and mycotoxin production (Figure 1, S2). The effect of lichen extract was assessed using the *Aspergillus parasiticus* mutant strain B62 (ATCC 24690) (Lee *et al.* 1971). *Aspergillus parasiticus* B62 accumulates the first stable aflatoxin precursor, norsolorinic acid (NA), due to a mutation in nor-1, and, due to biochemical redundancy, also accumulates reduced quantities of aflatoxins B_1_, B_2_, G_1_ and G_2_. Accumulation of NA, which is orange-red, offers a rapid visual screen to evaluate the activity of extracts on aflatoxin accumulation. To test the activity of lichen extracts, 10 μl, 30 μl, or 60 μl extract prepared as described above was added to 1 ml Glucose Mineral Salts (GMS), a chemically defined medium that induces aflatoxin, prepared as described by (Buchanan and Lewis (1984) and supplemented with 5 μM Zn_2+_. One ml GMS agar medium was added per well in a 24-well plate (Corning ®Costar, Sigma-Aldrich, St. Louis, MO, USA), and amended with lichen extract for the test wells or the corresponding volume of solvent to the control wells. After the medium was solidified, wells were center-inoculated with 2 μl of B62 spore suspension (10^5^ spores/ml) onto the surface of the medium and cultures were incubated in the dark at 29°C. To exclude position effects, wells with the extracts were compared with the asymmetrically positioned control wells in the same plate. Cultures were initially screened visually (phenotypic screening). Semi-quantitative assessments of aflatoxins and NA were performed using thin layer chromatography (TLC). TLC analysis was performed using plates coated with silica gel matrix containing a fluorescent indicator (#99876; Sigma-Aldrich, St. Louis, MO, USA) and run in the solvent system: chloroform:acetone=95:5. In addition, accumulation of NA was measured by direct scanning of the bottom surface of each well with a colorimeter Minolta CR-300 Chroma Meter (Konica Minolta, Osaka, Japan) and recording the “a” coordinate of the red-green axis in “color units” (Roze *et al.* 2007). Fungal growth was estimated by colony diameter.

**Figure 1.**
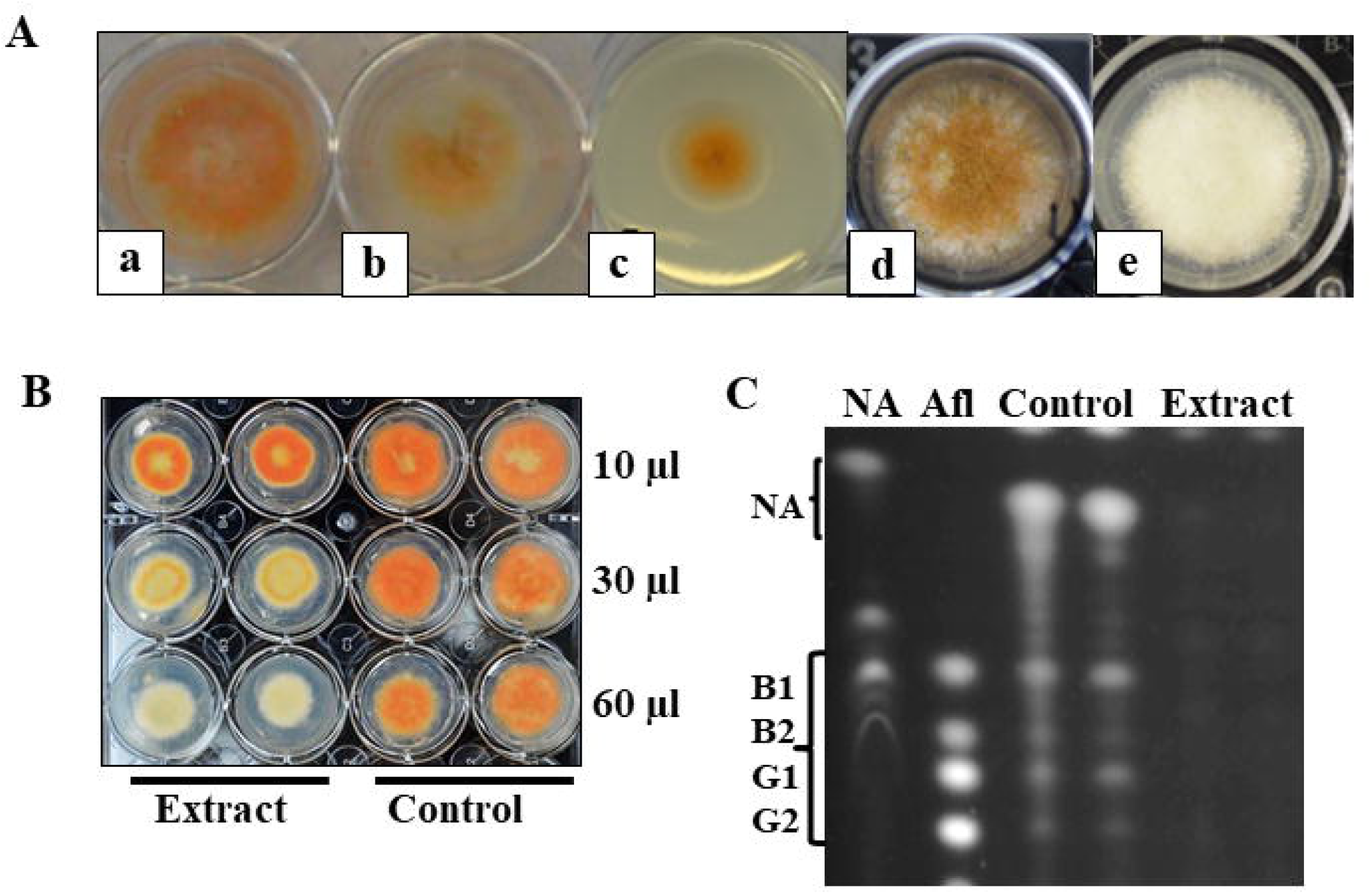
High through-put assay for detecting effects of lichen extracts on specialized metabolite production, growth and sporulation in *Aspergillus parasiticus* strain B62. Norsolorinic Acid (NA; orange-red pigment), accumulated by B62 is a precursor and indicator of aflatoxin production, used as a proxy for fungal specialized metabolite production. A, Morphological changes of treated cultures: (a) control (untreated) demonstrating normal growth and NA production compared with treated cultures exhibiting (b) NA biosynthesis inhibition; and (c) growth inhibition. (d) Control culture showing normal production of brown spores on the surface; (e) growth in presence of sporulation inhibitor. (a-c) Three-day old cultures on aflatoxin inducing medium and (d,e) 5-day old cultures on growth medium without aflatoxin induction. View from Above; B, Effects of dose of ethanol extracts from *Evernia prunastri* after 5-day exposure in culture. Numbers on the right indicate volume ethanol extract per 1 ml medium per well; control wells contain corresponding volumes of ethanol. View from Below; C, TLC analysis of aflatoxins extracted from cultures grown in aflatoxin inducing medium in the presence of *E. prunastri* ethanol extract (60 μl per 1 ml medium per well) for 5 days, aflatoxins B1, B2, G1, G2, NA. To view the entire TLC plate, showing the distortion of the NA lane, see Figure S1.

### Thin layer chromatography for assessment of effects on norsolorinic acid accumulation

To assess the inhibitory effects of lichen extracts on the accumulation of NA, cultures from treated and control cell wells were extracted with chloroform twice followed by acetone twice. Extracts were combined and evaporated and resuspended in 70% methanol to 1 ml. Culture extracts (5 μl sample per lane) and known quantities of aflatoxin standards (Sigma) were spotted on Partisils LHPKD silica gel TLC plates (60Å, 10 × 10 cm, 200 μm thick, plates; Whatman Inc., Clifton, NJ). Plates were resolved using a 95% chloroform in acetone (v/v) solvent system and analyzed under UV light. TLC plates were photographed using a Kodak DC 290 Zoom Digital Camera and band intensity was calculated using Kodak 1D 3.6 Image Analysis software. Band intensity and Rf values for bands were compared with values for the aflatoxin standards to establish relative concentrations and identity. Background intensity was subtracted from each band to obtain a true measure of band intensity (Roze *et al.* 2007).

### Analysis and quantification of trichothecenes

Wild-type *Fusarium graminearum* PH-1 (FGSC 9075; NRRL 31084)(Trail and Common 2000) was cultured in a medium that stimulates trichothecene production as previously described (Brown *et al.* 2004). Briefly, cultures were initially grown in liquid medium to generate mycelia. The mycelium from the culture was harvested, gently ground using a sterile mortar and pestle, and homogenized in a sterile glass homogenizer with a pestle. Homogenized mycelium was added to the trichothecene inducing medium. The suspension was mixed and aliquoted to 24-well plates. The wells were amended with 20 μl or 50 μl of lichen extract per well prepared as described above. Control wells contained corresponding volume of acetone or ethanol. After 6 days of incubation in the dark at RT contents of the wells were collected and lyophilized. Quantification of deoxynivalenol (DON), and 15-acetyl-deoxynivalenol (15ADON) in samples was performed in the Dong lab in the Department of Plant Pathology, University of Minnesota, St. Paul using gas chromatography (GC)-MS following the methods previously described (Mirocha *et al.* 1998).

### Liquid Chromatography-Mass Spectrometry/Mass Spectrometry

Liquid Chromatography-Mass Spectrometry/Mass Spectrometry (LC/MS/MS) was used to elucidate the profile (the presence and numbers) of compounds in the crude extracts and in the active fractions collected from flash chromatography (see below). Analyses were performed at the MSU Mass Spectrometry Facility. LC/MS/MS was performed as previously described (Staples *et al.* 2019). Briefly, to prepare samples for profiling of small molecules by LC-MS/MS, dried thallus from lichens was cut into small pieces (from 1 mm ×1 mm to 1 mm × 2 mm) and extracted twice with 2 ml ethanol or acetone. The combined extracts were dried under N_2_ and the pellet re-dissolved in 400 μl of the corresponding solvent. The solution was analyzed by LC-MS/MS (XEVO G2-XSQ TOF LC-MS/MS ESI, Waters, Milford, MA, USA) with 2 μl of extract injected into an Acquity HPLC CSH C18 column, (100 × 2.1 mm, 1.7 μM; Waters Corporation, Milford, MA, USA). The flow rate was maintained at 0.4 ml/min and the temperature was kept at 55°C. Two mobile phases were used. Mobile phase A consisted of acetonitrile (CH_3_CN:H_2_O)/60:40 + 10 mM ammonium formate + 0.1% formic acid. Mobile phase B consisted of CH_3_CN:H_2_O/90:10 + 10 mM ammonium formate +0.1% formic acid. A linear gradient elution program was applied as follows: 0 min, 40 % B; 0–2 min, 43 % B; 2–2.1 min, 43–50 % B; 2.1–12 min, 50–54 % B; 12–12.1 min, 54–70 % B; 12.1–18 min, 70-99 % B; 18–18.1 min 99-40% B; and 18.1–20 min 40% B. MS data were recorded in negative ionization mode using MassLynx 4.1. software (Waters, Milford, MA, USA). All solvents were HPLC grade (Sigma-Aldrich, St. Louis, MO, USA).

### Flash Chromatography

Flash chromatography was used as part of the purification of methylorsellinate, an active compound in *Physcia stellaris* (Figure 2, S3). Active fractions were then further purified by HPLC and analized with LC/MS/MS. Briefly, 2 g of dried lichen were prepared as above and extracted twice with 10 ml 100% ethanol. The extracts were combined, centrifuged at 10,000 rpm for 20 min and the supernatant applied to a silica column (pore size 60 Å, particle size 230-400 mesh which equals 40-63 μm; column diameter 18 mm, length 40 cm, silica gel height in column 20 cm, and silica volume in the column 50.8 cubic centimeters; Fluka, Sigma-Aldrich, St. Louis, MO, USA) that was equilibrated in hexane. The column was eluted with hexane:ethyl acetate (1:0–1:3) collected in 11 fractions, followed by hexane:ethyl acetate:acetone (1:4:1), hexane:ethyl acetate:acetone (1:5:2), and finally with ethyl acetate:acetone (5:3). Each of the total 20 fractions consisted of 15 ml. The fractions were evaporated under N_2_, and re-dissolved in 0.5 ml ethanol. Activity of each fraction was assayed for effect on spore germination and NA accumulation (Figures 2, 3, S3).

**Figure 2.**
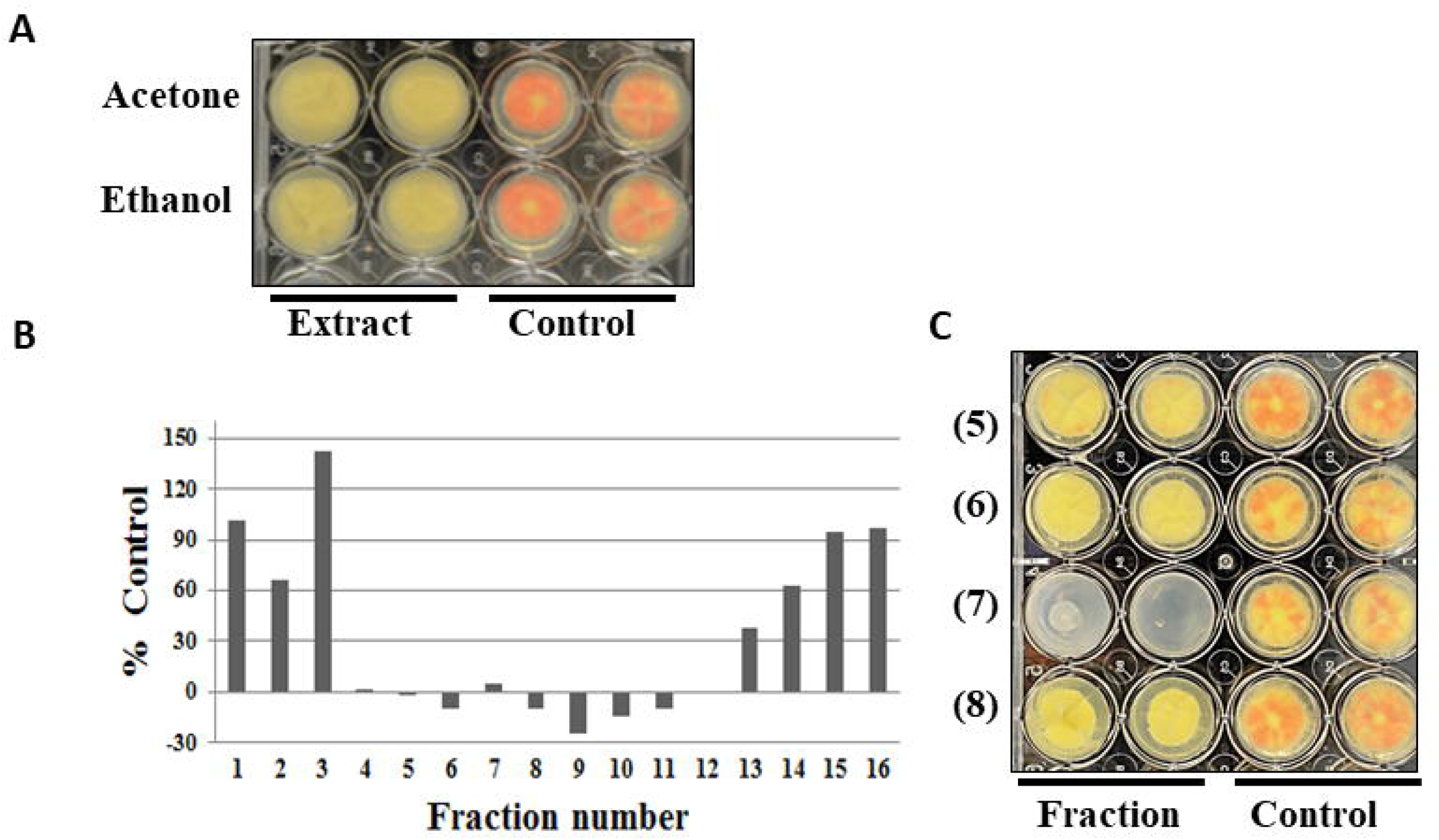
Purification and characterization of an inhibitor of norsolorinic acid biosynthesis, methylorsellinate, from *Physcia stellaris*. A, *P. stellaris* holobiont ethanol and acetone extracts bioassay was performed as described in Methods using 60 μl of extracts. View from below. B. Inhibition of NA accumulation by flash chromatography fractions. Intensity of NA was measured by the ChromaMeter; negative values reflect absorption values in the reading. Note that Fraction 7 inhibits growth; C,Bioassay of flash chromatography fractions; fraction number indicated. Control is amended with ethanol. NA is low in the controls compared to some assays, but within the normal range.

**Figure 3.**
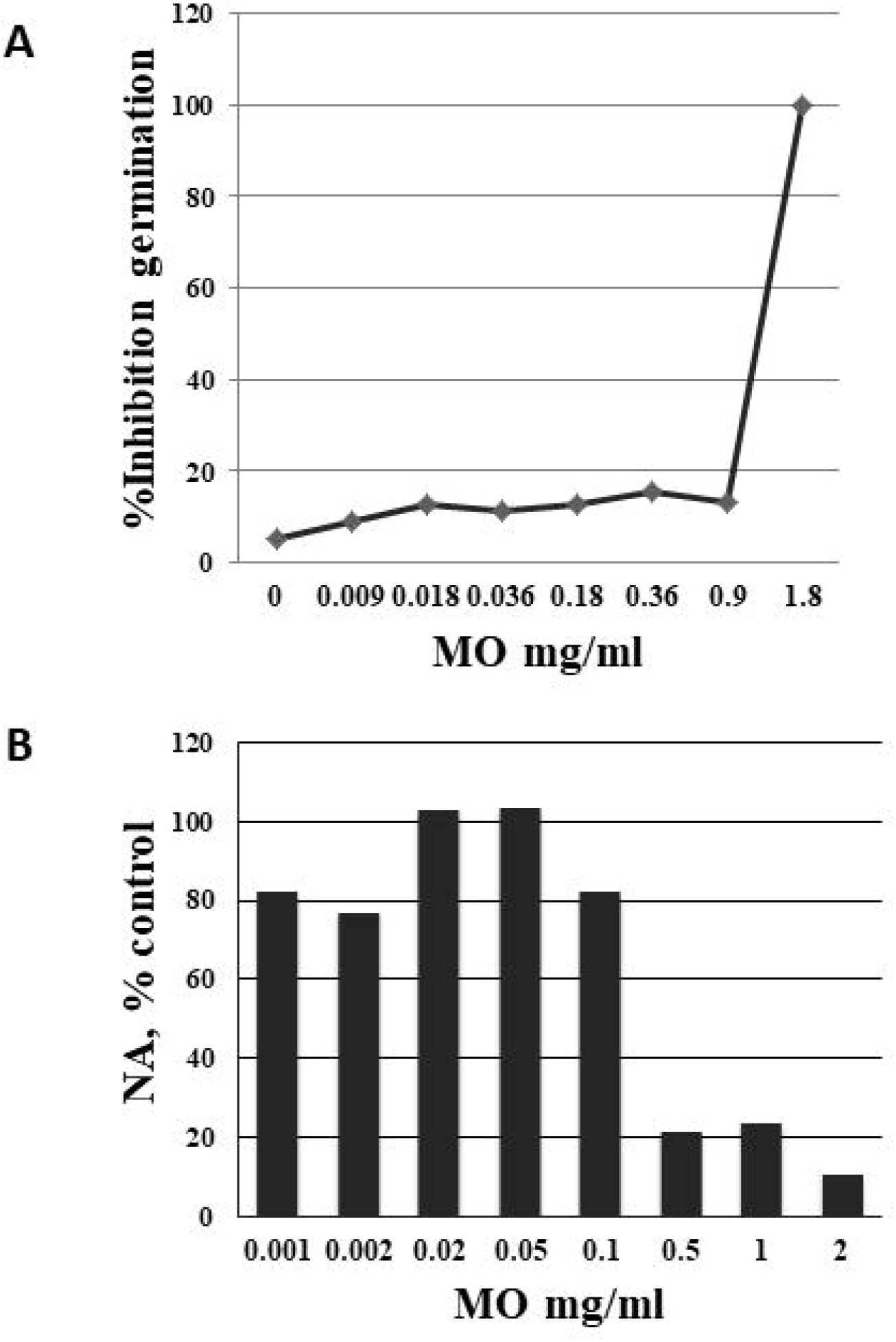
Effects of methylorsellinate (MO) on norsolorinic acid accumulation and germination in *Aspergillus parasiticus*. MO demonstrates dual activity: it inhibits germination of *A. parasiticus* and inhibits NA/aflatoxin accumulation. Spore germination assay was carried out on a microscope slide in drops of liquid GMS medium placed in a moist chamber. Numbers of germinated and ungerminated spores from three randomly selected areas per slide were counted and the spore germination percentage calculated.

### The spore germination assay

The assay was carried out on a microscope slide in drops of liquid GMS medium placed in a moist chamber. Numbers of germinated and ungerminated spores (conidia) from three randomly selected areas per slide were counted and the spore germination percentage calculated as a ratio of non-germinated spores per total number of spores multiplied by 100%.

### High Performance Liquid Chromatography (HPLC) analysis

The HPLC analysis was performed as described by Feige *et al.* (1993) with modifications described by (Staples *et al.* 2019). Briefly, a Spherisorb 5 ODS2 silica-based, reversed-phase C18 column (5 μm, 250 × 4.6 mm; Waters Corporation, Milford, MA, USA) was used. The column was operated at room temperature. Kontron HPLC system with data system 450 instrument, 430 UV detector and 360 autosampler were employed. Compounds were detected at 245 nm and the UV spectra (200–400 nm) of each peak eluted were recorded automatically. Solvent system: A - Deionized Milli-Q water containing 1.0% orthophosphoric acid, B −100% methanol (J.T. Baker Chemical Co, Phillipsburg, NJ, USA). Gradient: run started with 30% B and continued isocratically for 1 minute at flow rate 0.7 ml/min. After 1 minute 20 μl of sample was injected. System B was increased to 70% within 14 minutes and then to 100% within the next 30 minutes. Run was continued isocratically at 100% B for further 18 minutes (total run time 62 minutes). At the end of the run, solvent system B was decreased to 30% within 1 minute and column flushed with 30% B for 16 minutes before a new run was initiated. The pipeline for discovery of bioactive lichen compounds is summarized in Figure S1.

### Microbiome analysis

*Cladonia cristatella* (81b)*, C. rangiferina* (117b), and *Evernia mesomorpha* (75), and *Physcia stellaris* (82b) were used for microbiome sampling. After collection, samples were brushed free of debris, but were not surface sterilized because it was difficult to avoid absorption of the decontamination solution by the delicate and convoluted lichen thallus. Therefore, results include the epiphytic and endophytic microbes. The samples were lyophilized and stored at −80°C until processing. DNA extraction was performed using a NucleoSpin Soil Kit (Takara Bio USA, Inc., Mountain View, CA, USA) according to the manufacturer’s instructions with the following optimizations; all lyophilized samples were ground with a sterile mortar and pestle before applying to the NucleoSpin Bead Tube (Type A) containing ceramic beads and Buffer SL2; no Enhancer SX was used.

DNA from microbial communities was amplified using primers for the ribosomal ITS2 region used to identify fungi (Toju *et al.* 2012) and oomycetes (Cooke *et al.* 2000; Sapkota *et al.* 2015), and the v4 region of the 16S rRNA gene to identify bacteria (Kozich *et al.* 2013). PCR amplification of each sample was performed in triplicate, PCR products were pooled and purified with Wizard Gel and PCR Clean-up Kit (Promega, Madison, Wisconsin, USA). Purified PCR products were sequenced at Michigan State University. Demultiplexed sequences were analyzed using the USEARCH pipeline version 8.1. Bacterial OTUs were classified using the RDP 16S database. Fungal OTUs were classified using the RDP ITS database. A custom oomycete database was created using sequences from Robideau *et al.* (2011) trimmed to the ITS2 region. Sequences have been deposited in the NCBI SRA (accession number pending). Microbial communities from each organism group were analyzed in parallel. Taxa with fewer than 5 total reads across all samples were removed. To determine the microbial community diversity within lichens, comparisons were made between the four host species. Diversity indices were calculated using the *phyloseq* package (McMurdie & Holmes 2013) in R. Alpha diversity was determined by Shannon Index (H’), and differences between H’ of each host species were determined with ANOVA and Tukey’s HSD. Ordination analyses were conducted with Non-metric multidimensional scaling (NMDS) of the Bray-Curtis index, and ANOSIM was used to determine differences in microbial communities of each lichen species.

## Results

### Lichen extracts affect fungal specialized metabolite production, sporulation and hyphal growth

To identify the presence and frequency of three major effects lichens can have on fungi in their microbiomes (radial growth, mycotoxin production and spore production), we developed assays based on two well-studied fungi that produce mycotoxins: *Aspergillus parasiticus* and *Fusarium graminearum*. Mycotoxins are easily quantified as they are routinely evaluated in food and feed due to their deleterious effects on human and animal health. In the screens, we used *A. parasiticus* strain B62, which accumulates orange-red norsolorinic acid (NA), a precursor and indicator of aflatoxin production, under aflatoxin inducing conditions (Figure 1). All lichens were tested through the activity pipeline (Figure S1). Initially, ethanol, acetone, hexane or ethyl acetate were used as solvents, but hexane and ethyl acetate extracts generated effects in only 8 of 17 lichens tested and lichen extracts did not affect all phenotypes (Table 1; Table S1). Ethanol and acetone extracts were found to be the most effective solvents for extraction of the desired activities, and we focused further efforts on these (Table S1).

**Table 1.**
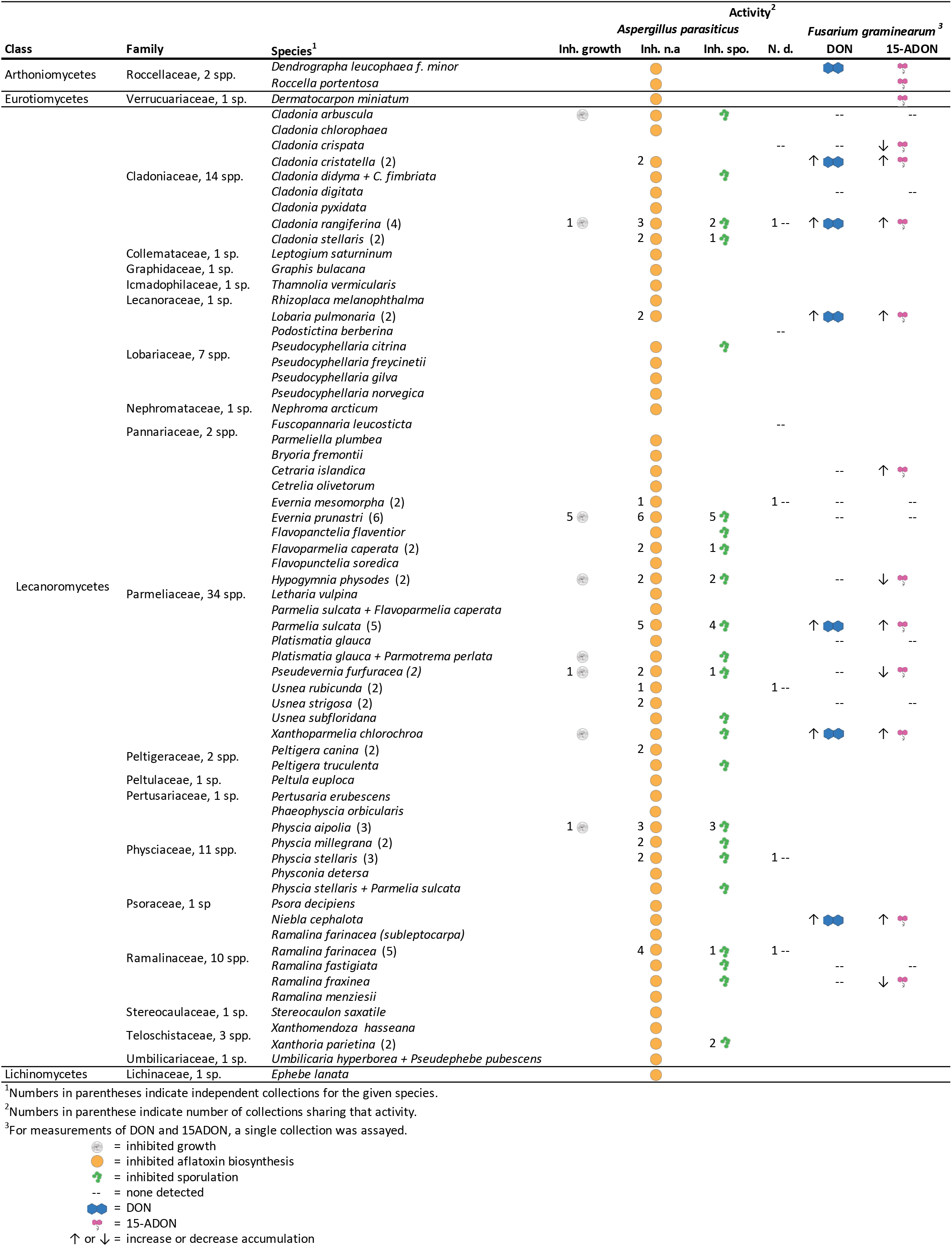
A phylogenetic survey of antifungal activities in lichen extract. Activities of species tested within each family are represented as designated in the key. Each symbol represents the activity in one lichen, some lichens may have more than one activity with *Aspergillus parasiticus* as the model fungus.

To expand our understanding of activity in lichen extracts on accumulation of specialized metabolites in fungi, we tested their effect in *F. graminearum* on biosynthesis of the following mycotoxins: the carmine polyketide aurofusarin, and trichothecene mycotoxins deoxynivalenol (DON) and 15-acetyl deoxynivalenol (15-ADON). Out of 21 extracts tested twelve lichen extracts reduced DON accumulation to undetectable levels, while twelve reduced levels of the 15-ADON derivative, some of these affected production of both trichothecene mycotoxins (**Table S2**). Surprisingly, extracts of eight lichens increased DON and ten extracts increased 15- ADON, effects we did not see in the assays on NA accumulation.

Ethanol extracts from the 67 lichen species caused cultures to exhibit: inhibition of NA accumulation (62 species); reduced hyphal growth (8 species); decreased (19 species) and activated (2 species) spore production. Six species did not show any activity and several species exhibited more than one activity (**Tables 1, S1**). In summary, inhibition of specialized metabolite production, as determined by our visual assay, is the most frequently observed activity, being present in 92% of the 67 lichen species analyzed across the phylogeny.

### Purification, identification and activity of methylorsellinate from Physcia stellaris and effect on fungi

The crude ethanol extract prepared from *P. stellaris* thalli was separated by flash chromatography and all 20 fractions were screened by high through-put activity assay for effects on mycotoxin biosynthesis, hyphal growth and sporulation. Fraction 7 demonstrated significant inhibition of fungal growth, NA production and sporulation. Fraction 7 was analyzed by HPLC and LC-MS (Figures 2, S3). The most abundant ion in negative mode had a mass of m/z 181.05, the exact mass for methylorsellinate (Musharraf *et al.* 2015 Anal Methods). Purified methylorsellinate, depending on concentration, demonstrated dual activity with effects on spore germination and accumulation of NA (Figure 3). Based on retention times (RT) of 20.2–20.3 and the spectra we suggest that this compound is present at various levels in other lichens we tested as well: *E. prunastri*, *Hypogymnia physodes*, *Parmelia sulcata*, *Platismatia glauca*, *Pseudevernia furfuracea* (Data not shown).

### *Culture of the lichenized fungus and alga of* Cladonia cristatella *and activity of holobiont vs. mycobiont*

To investigate the role of mycobionts in producing the active compounds, we grew mycobionts and photobionts of eight lichens in culture. Fungal cultures were grown on MEA from isolated ascospores obtained from *C. cristatella*, *L. pulmonaria*, *Physcia aipolia, P. millegrana*, *P. furfuracea*, *R. fastigiata*, *R. fraxinea*, and *Xanthoria parietina*. Interestingly, we found that extracts from the mycobiont culture of *C. cristatella* (Figure 4), as well the lichen holobiont, strongly inhibited specialized metabolism (NA production) of *A. parasiticus* (Figure 5). However, we were not able to determine the active compound/s which possesses the activity since many unidentified compounds were present in extracts despite the attempts to purify and concentrate active compounds.

**Figure 4.**
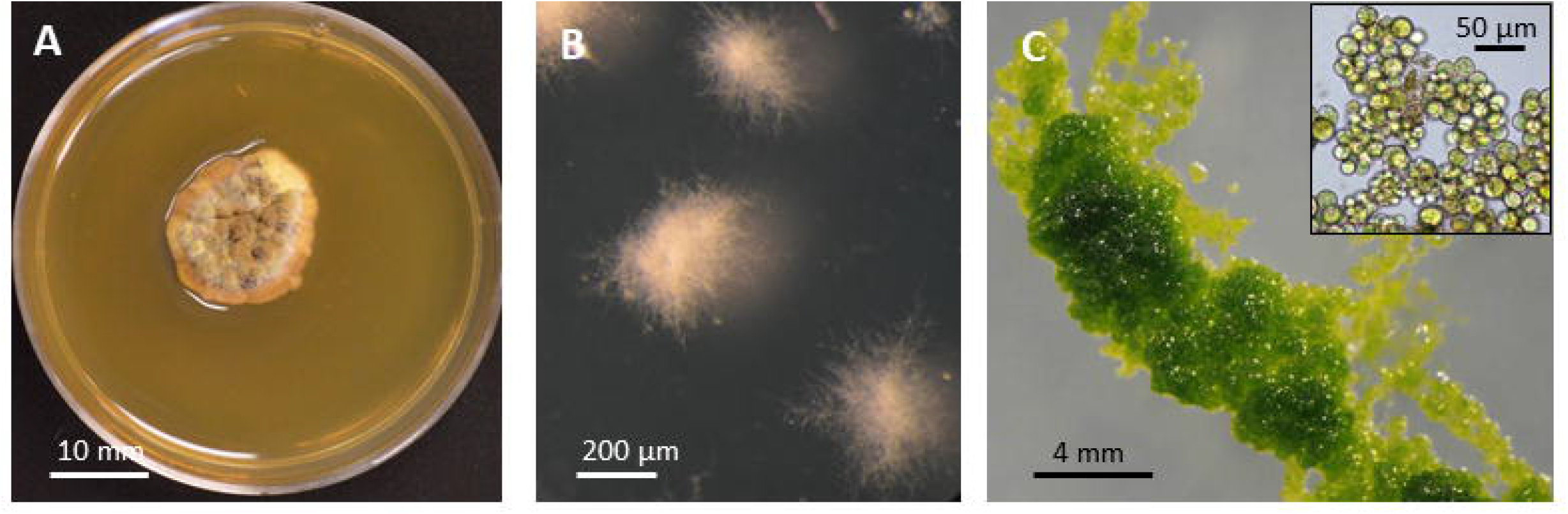
Mycobiont and photobiont of *Cladonia cristatella* in culture. A, Mycobiont in culture grown on solid medium for two months; B, Mycobiont in submerged culture; C, Photobiont grown on solid medium, 4 week old culture. Inset: Photobiont in liquid medium.

**Figure 5.**
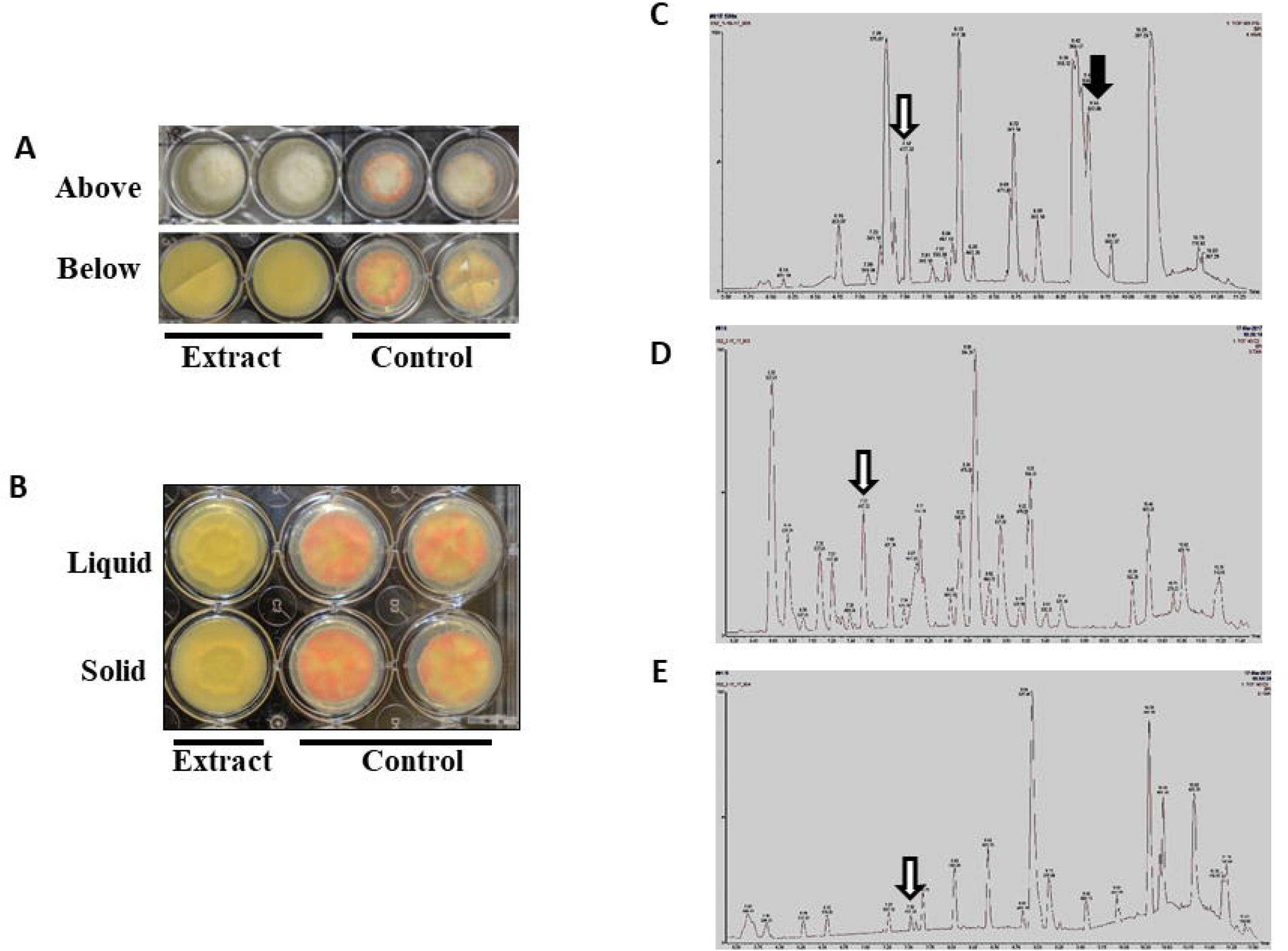
Extracts of liquid and solid cultures of *Cladonia cristatella* mycobiont lower accumulation of norsolorinic acid (NA). A, Holobiont ethanol extract bioassay. View from above and below; B, Bioassay of extract of mycobiont was grown in liquid and solid malt extract (ME) medium for 5 weeks, extracted with ethanol and partially purified and concentrated using Solid Phase Extraction cartridges (SPE, OASIS, Waters, USA); view from below. C-E, LC-MS chromatograms of ethanol extracts showing reduction in NA accumulation; Unknown compound (White arrow in all three samples; ionic mass 417.32 at 7.52) was present in all 3 extracts analyzed and is a candidate for active compound; C, crude ethanol extract. Usnic acid (Black arrow; ion mass 343.08 at 9.55); D, ethanol extract of mycobiont solid culture after SPE recovery; E. ethanol extract of mycobiont liquid culture after SPE recovery. Bioassays of A and B were performed as described in Methods using 60 μl of respective extracts.

### Lichen Microbiome

Three lineages of microorganisms (bacteria, fungi, and oomycetes) were identified as additional members of the microbiomes (Figure 6). Four lichen species were sampled for the microbiome study as they represent three lichen habits: *Cladonia cristatella* and *C. rangiferina* (*Cladoniaceae*) are dimorphic fruticose species that are predominantly found on the soil surface; *E. mesomorpha (Parmeliaceae)* is a loosely attached fruticose species and often inhabits branches of trees and similar habitats; and *P. stellaris (Physciaceae*) is a closely attached foliose species and frequently inhabits aerial woody plant parts. Richness of fungal and bacterial communities differed by host lichen (p < 0.05; Figure 6A, 6D) but not oomycetes (Figure 6G) and the community composition of each microorganism lineage also differed by host lichen (p < 0.05, Figure 6B, 6E, 6H). The fungal members of the microbiome were, as expected, dominated by the lichenized fungal species of each lichen (Figure 6C) but species of Dothidomycetes and Eurotiomycetes were the dominant additional fungal members of the microbiome of *E. mesomorpha, P. stellaris,* and *C. rangiferina* (Figure 6C, Table S). *Cladonia cristatella* harbors those classes, but is different in having a notable presence of the chytrid *Powellomyces* sp., commonly associated with soil and decaying plant tissue (Simmons 2011) and member(s) or the basidiomycete order *Tremellales* which is common on rotted wood, exactly where this lichen was collected.

**Figure 6.**
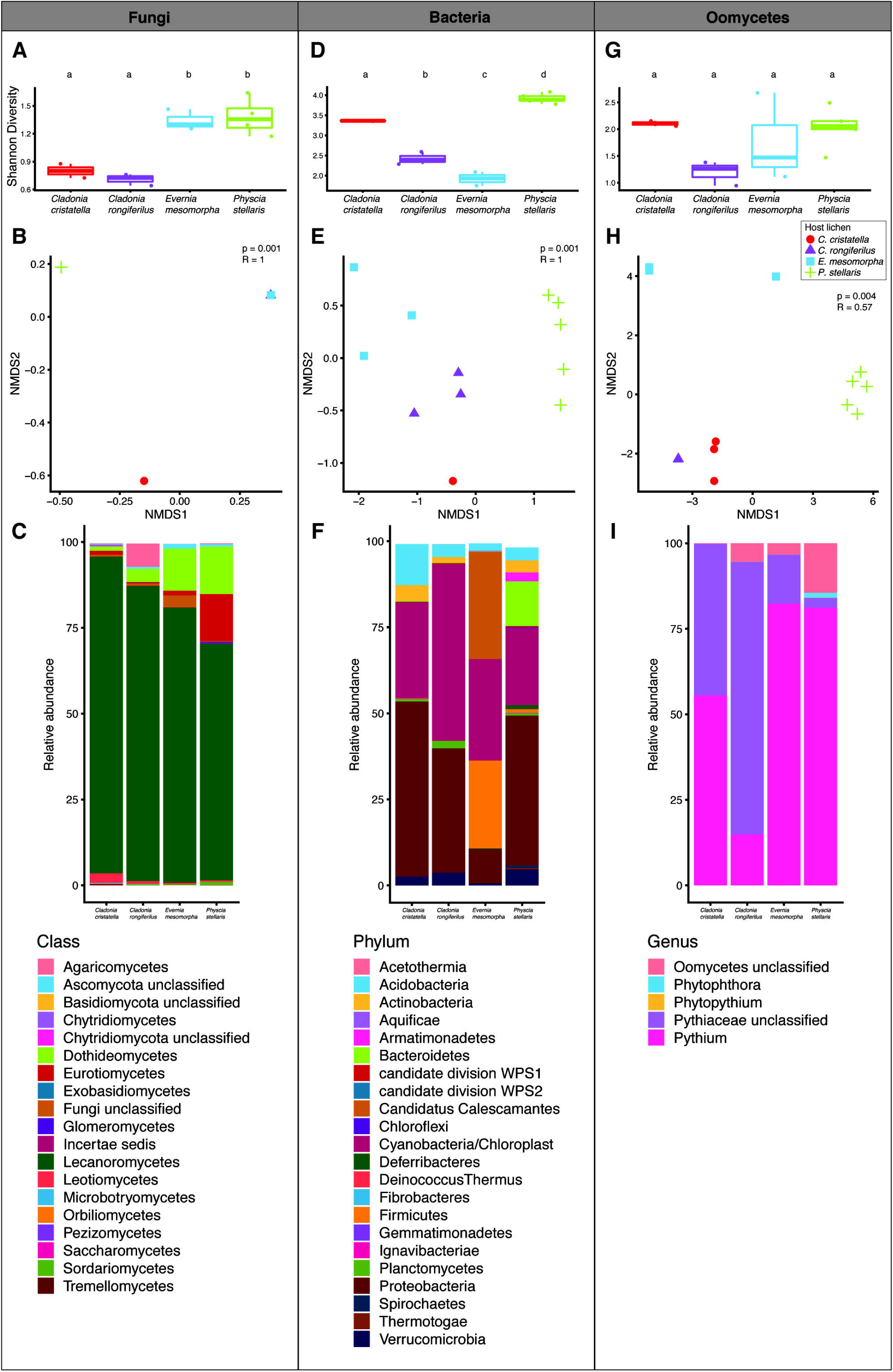
Diversity of lichen microbiome. Shannon diversity index (H’) of fungi. (A) bacteria (D), and oomycetes (G) members of lichens; letters indicate pairwise comparisons with p < 0.05 after ANOVA and Tukey’s HSD. Non-metric multidimensional scaling of Bray-Curtis Index of fungi (B), bacteria (E), and oomycetes (H); p and R values from ANOSIM between host lichen species. Points in each panel represent calculated index from replicate samples, colored by host lichen species. Relative abundance for fungi (C), bacteria (F), and oomycetes (I) in each host lichen.

Comparisons of bacterial communities between the lichen hosts demonstrated a predominance of Cyanobacteria/chloroplasts (from the photobiont) and Proteobacteria. *H’* differed by lichen host species (p < 0.05, Figure 6D). *Physcia stellaris* was host to Actinobacteria, Armatimonadetes, and Bacteroidetes species, which were present at negligible levels in the other lichen species, and *P. stellaris* had the highest *H’* for both bacteria and fungi (Figure 6F). *Evernia mesomorpha* showed a divergent pattern of colonization with *Candidatus calescamantes* (Becraft *et al.* 2016) and Firmicutes as dominant taxa, two lineages containing many anaerobic bacteria, in addition to Cyanobacteria and Proteobacteria (Figure 6F). Firmicutes are common microbes in animal guts but have been preliminarily suggested to be associated with the microbiota of fungi (Paul *et al.* 2019a; 2019b). The oomycete communities were dominated by Phytophthora, Pythium, members of the Pythiaceae and unclassified oomycetes (Figure 6I). Members of the Pythiaceae were present in all host lichens but were the dominant taxa in *Cladonia* spp. In summary, in overall community composition, each microorganism lineage differed by host lichen (p < 0.05, Figure 6B, 6E, 6H).

## Discussion

Elucidating the role of specialized metabolites in the mechanisms underlying lichen assembly and function has important implications for understanding not only lichen community assembly but for revealing the fundamental processes in microbiota in general. The consistent presence of activities that manipulate fungal specialized metabolism across the phylogenetic tree indicates this is an important aspect of lichen integrity. Members of the lichen community may manipulate specialized metabolism of the essential and transient fungal members and thus attenuate negative interactions with the incumbent fungi or support the beneficial fungal partners.

The majority of lichens tested here exhibit regulatory activity towards specialized metabolism of *F. graminearum* and *A. parasiticus.* In contrast, very few lichen extracts were active in regard to fungal sporulation (14%) and growth (4%). These results appear counterintuitive, as the host lichen would seemingly want to limit the biomass and reproductive abilities of the endophytic microbes. However, regulation of sporulation appears to be fairly conserved as a fungal developmental process (Lee *et al.* 2016) as is hyphal growth (reviewed by Steinberg *et al.* 2017). If the ability to control the microbiome by chemicals specific can be effective and discerning, allowing the microbiome members to be controlled so they can be beneficial without being invasive, then the strategy would be effective and energetically thrifty. As a caveat, we tested only two fungal species, and therefore have limited data on which to speculate. Very few lichens lacked all of the activities, and the activities affecting specialized metabolism were distributed across the lichen phylogeny, indicating possible retention of these activities from a common ancestor or selection of these activities due to their beneficial presence.

We have demonstrated that methylorsellinate (MO) decreases accumulation of norsolorinic acid (NA) and aflatoxin at similar concentrations, and inhibits spore germination, sporulation and fungal growth in *A. parasiticus*. MO is a derivative of orsellinic acid and is formed through the acetyl-polymalonyl pathway, one of the three major biosynthetic pathways for specialized metabolites in lichens (Calcott *et al.* 2018). Orsellinic acid is one of the basic units in the biosynthesis of lichen phenolics (Gaucher & Shepherd 1968). Lecanoric acid is composed of two molecules of orsellinate (Huneck & Yoshimura 1996) and we have shown that lecanoric acid affects the aflatoxin pathway, and inhibits sporulation. Lichen phenolic compounds are mainly depsides, depsidones, dibenzenofurans (Calcott *et al.* 2018). The depsides are a large group of unique lichen polyphenolic compounds composed of two or more monocyclic units of orsellinic acid derivatives. Lichen phenolics are generally secreted by the fungal partner and deposited as crystals on the surface of the cell wall of the fungal hyphae (Hyvärinen *et al.* 2000). Although we have characterized the active components of the activity in these few cases, we cannot make a judgement on where the activities originate in the vast majority of our assays. It is important to look at these activities as outcomes of the whole community that are likely useful to its function.

Extracts from eleven lichens, *C. arbuscula, C. crispata, C. digitata, E. mesomorpha, E. prunastri, H. physodes, Platismatia glauca, Pseudevernia furfuracea, Ramalina fastigiata, R. fraxinea,* and *Usnea strigosa,* blocked accumulation of DON and its biosynthetic precursor 15-ADON. Interestingly, extracts of eight lichens tested increased the production of the trichothecene mycotoxins, which was an effect not observed in the assays on aflatoxins. DON and derivatives are produced when *F. graminearum* is infecting plants (Jansen *et al.* 2005) and is a virulence factor (Proctor *et al.* 1996). An interesting hypothesis is that the increase in DON reflects a similar mechanism when the fungus encounters the lichen as when it encounters the host plant. Complex cellular components and physiological factors such as pH, nutrients, light, genetic regulation and signaling pathways have been shown to affect DON production in the fungus. These have been recently comprehensively reviewed (Gardiner *et al.* 2009).

We have begun the process of purifying and characterizing active components in the lichens that induce and suppress trichothecene accumulation. Untangling components of lichen extracts that affect the production of trichothecenes will have meaningful consequences for agriculture and food safety. The regulatory activity preserved in herbarium specimens that we assayed indicates that lichens have the ability to preserve specialized metabolites for a very long time, which makes sense in light of their longevity and colonization of extreme environments. These tendencies are conserved in that lichens of different families also preserve mycotoxins for decades (Burkin & Kononenko 2015; Burkin *et al.* 2012). Several mycotoxin-producing fungi, including *Alternaria, Aspergillus, Fusarium,* and *Penicillum* are common members of the lichen microbiome, some of which were recorded here. Aflatoxin is a potent mutagen (Benkerroun, 2020), deoxynivalenol is an effective inhibitor of protein biosynthesis and causes apoptosis (Shifrin and Anderson, 1999), and zearalenone can affect developmental processes in animals and fungi (Calvo *et al.*, 2002). Hence lichens require a means to protect its microbiome not only from the adverse effects of mycotoxins and broad inhibition is an efficient method of doing this.

The effects of environmental specialized metabolites on fungi has been shown to affect morphological development and resistance to environmental stresses (Calvo *et al.*, 2002; Brakhage 2013). For example, volatile compounds may affect asexual sporulation, sclerotia formation, pigment and toxin production in fungi (Roze *et al.* 2007; 2011). An additional example demonstrates that the ability to produce a toxic secondary metabolite aflatoxin by Aspergilli may enhance their resistance to oxidative stress in comparison to atoxigenic strains (Roze *et al.* 2015). DON has been associated with virulence in wheat, gliotoxin is linked to *A. fumigatus* virulence during the infection process in humans (Scharf *et al.* 2016). These studies enhance a large body of research indicating that the ability to manipulate specialized metabolism in a fungal cell, specifically to inhibit or activate it, may serve as a powerful tool to weaken or enhance specific characteristics of a fungus.

In some lichen species, extracts from different collections varied in activity. These inconsistencies could be due to differences in the microbiome, or associated with growth stage, nutritional status, environmental factors, etc. LePogam *et al.* (2016) applied LDI-MS (Laser Desoprtion and Ionization-Mass Spectroscopy) for imaging the localization of specialized metabolites in lichen thalli. They demonstrated the sequestering of specialized metabolites in discrete locations in the thallus, depending on the metabolite. Certain specialized metabolites may be indicative of specific habitats and species composition in those habitats (Toni & Piercey-Normore 2013). A recent global study of distributions of lichens and associated endolichinic fungi showed that host selection drove the composition of lichen endophytes (U’Ren *et al.* 2019). The relationship between lichens, specialized metabolites, microbiome composition, and environment is very complex and part of this understanding the perceived variability of all of these factors and how they affect each other.

The microbiomes of four lichen species were characterized to provide some context for interpreting the extracted activities. Because the vast majority greatly reduced NA production, and increased or decreased trichothecene production, and there were fewer activities affecting sporulation and hyphal growth, it was not possible to make meaningful associations between types of lichens and their activities. Endolichenic fungi colonize healthy, photosynthetic tissues of lichens (U’Ren *et al.* 2019; U’Ren *et al.* 2012). The major endolichenic fungi were identified to species and for each of the 4 host species, their distributions showed a difference in colonization at higher taxa. The *Cladonia* spp. harbored fewer fungi and more bacteria, whereas *Evernia* showed the opposite pattern (Lawrey *et al.* 2003) Oomycetes were present across all species and to our knowledge they have not been identified previously as common components of lichen microbiomes. Fungal endophytes occur in lichens similar to their association with plants. They play important roles in lichen health, but the roles of these and the oomycetes are not well defined. Future experiments using metagenomics tools are essential to sort out roles. A recent large study of Dothidiomycetes demonstrates how analysis of large-scale sequencing can be used to clarify evolution of lifestyles (Haridas *et al.* 2020). The complex and multikingdom holobionts that are lichens, combined with their infinite variation, will require much greater facility with analysis of the results of such a study.

## Acknowledgements

We thank Dr. A. Schilmiller and the MSU Mass Spectrometry Facility for the mass spectrometry analyses. We greatly appreciate the analysis of DON metabolites by Dr. Yanhong Dong, Dept of Plant Pathology, Univ. of Minnesota. The authors also thank Dr. Zachary Noel for assistance in generating the Oomycete database. Funding was provided by a grant from the MSU Office of the Vice President of Research and Graduate Studies, AgBioResearch and the College of Natural Sciences.

---

**Table S1.** Lichen provenance and activities.

**Figure S1.** Pipeline for discovery of bioactive lichen compounds used in this paper.

**Figure S2.** Intact TLC plate from Figure 1C showing the NA band distortion and thus demonstrating that in lanes 3 and 4 the second band from the top lines up with the NA control in the first lane. 1:*Evernia prunastri.* 17: *Physcia aipolia*, (collection 18). 19: *Parmelia sulcata.* Afl, aflatoxin standards containing aflatoxins B1, B2, G1, and G2. NA, norsolorinic acid.

**Figure S3.** Chemical characterization of methylorsellinate acid from *Physcia stellaris.* A, Chromatogram of fraction 7 (from Figure 3). AU, absorbance units at 260 nm; B, Absorbance spectrum of fraction 7. Absorbance maxima at 268 and 304 nm.

