## Supplementary figures and images for "Activity throughout the lichen phylogeny indicates a focus on regulation of specialized metabolites"

### Figure S1

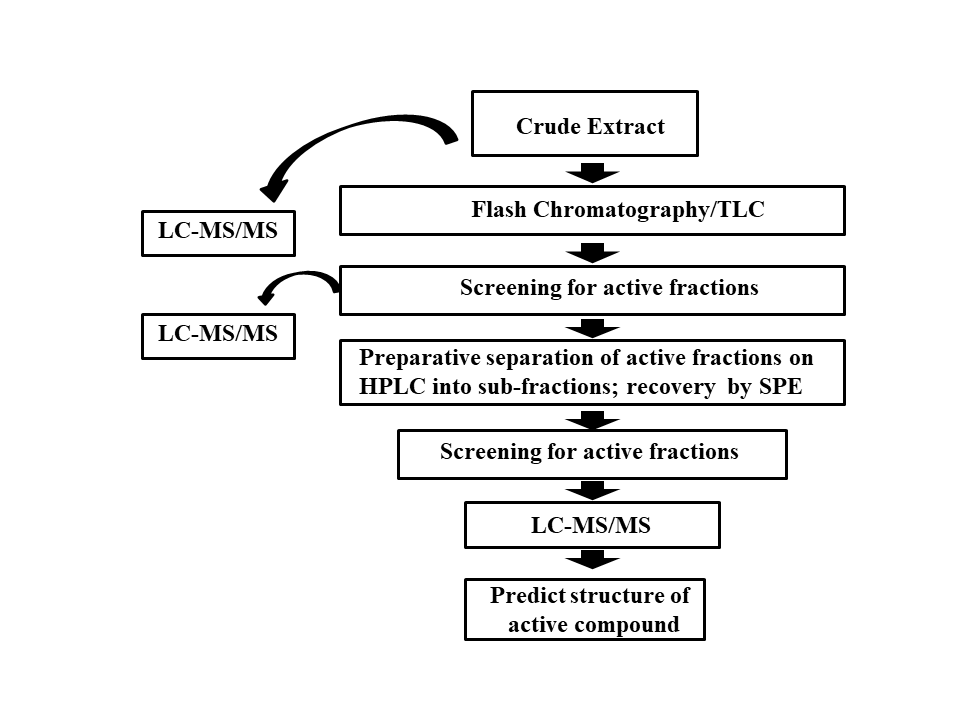

### Figure S2

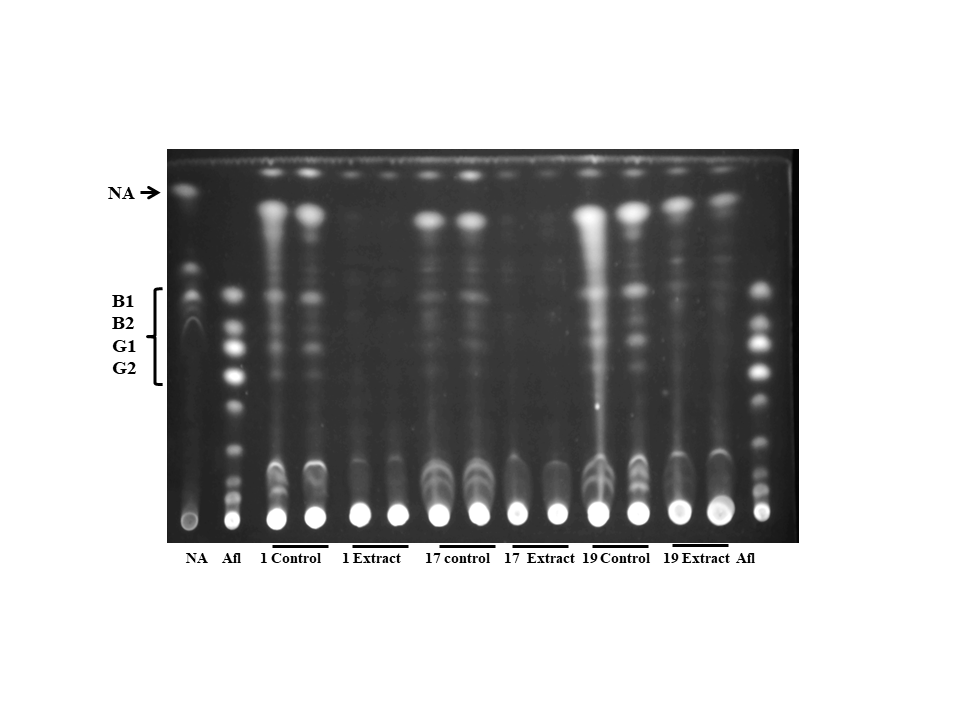

### Figure S3

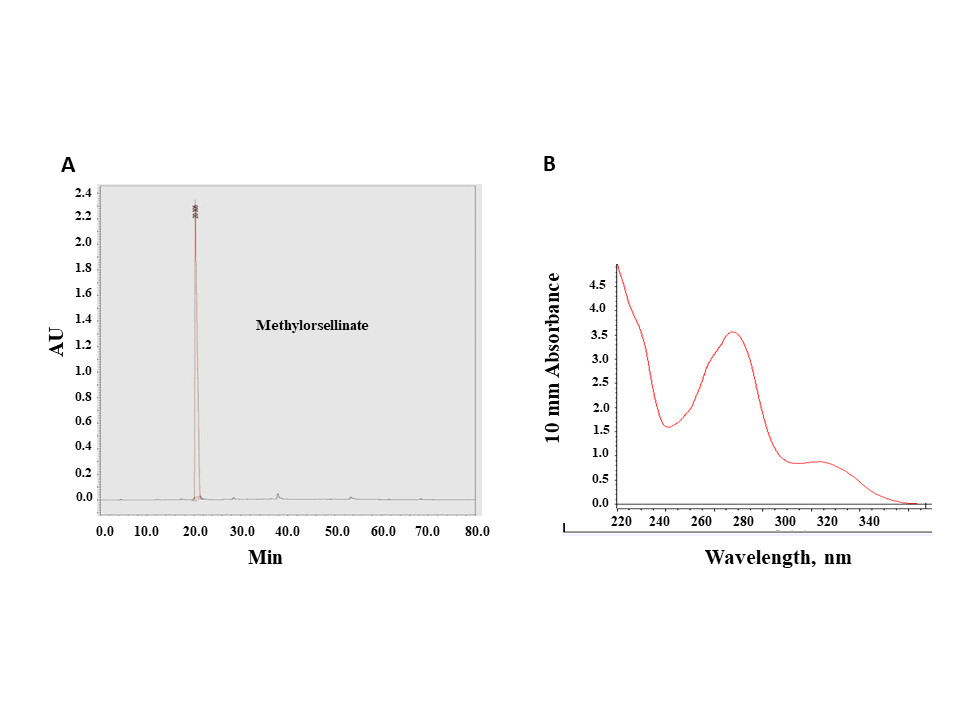
